# Genome wide assessment of genetic diversity and transcript variations in 17 accessions of the model diatom *Phaeodactylum tricornutum*

**DOI:** 10.1101/2023.05.31.543125

**Authors:** Chaumier Timothée, Feng Yang, Eric Manirakiza, Ouardia Ait-Mohamed, Yue Wu, Udita Chandola, Bruno Jesus, Gwenael Piganeau, Agnès Groisillier, Leila Tirichine

## Abstract

Diatoms, a prominent group of phytoplankton, have a significant impact on both the oceanic food chain and carbon sequestration, thereby playing a crucial role in regulating the climate. These highly diverse organisms show a wide geographic distribution across various latitudes. In addition to their ecological significance, diatoms represent a vital source of bioactive compounds that are widely used in biotechnology applications. In the present study, we investigated the genetic and transcriptomic diversity of 17 accessions of the model diatom *Phaeodactylum tricornutum* including those sampled a century ago as well as more recently collected accessions. The analysis of the data reveals a higher genetic diversity and the emergence of novel clades, indicating an increasing diversity within the *P. tricornutum* population structure, compared to the previous study and a persistent long-term balancing selection of genes in old and newly sampled accessions. However, the study did not establish a clear link between the year of sampling and genetic diversity, thereby, rejecting the hypothesis of loss of heterozygoty in cultured strains. Transcript analysis identified novel transcript including non-coding RNA and other categories of small RNA such as PiwiRNAs. Additionally, transcripts analysis using differential expression as well as Weighted Gene Correlation Network Analysis has provided evidence that the suppression or downregulation of genes cannot be solely attributed to loss of function mutations. This implies that other contributing factors, such as epigenetic modifications, may play a crucial role in regulating gene expression. Our study provides novel genetic resources, which are now accessible through the platform PhaeoEpiview (https://PhaeoEpiView.univ-nantes.fr), that offer both ease of use and advanced tools to further investigate microalgae biology and ecology, consequently enriching our current understanding of these organisms.

## Introduction

Photosynthetic microalgae are important components of life in the oceans providing organic biomass and fueling a range of key biogeochemical processes. Diatoms in particular are widely recognized as one of the most significant phylum of phytoplankton, owing to their substantial contribution to primary productivity, carbon fixation, and biogeochemical cycling of essential nutrients such as nitrogen and silicon [1, 2]. In addition to their ecological importance, diatoms are a rich source of bioactive compounds with diverse applications in various industries, including nutraceuticals, nanotechnology, pharmaceuticals, and food and feed industries [3, 4]. In recent years, using model species in diatoms has dramatically increased our knowledge about the biology and ecology of these important organisms [5]. Particularly, one species, the diatom *Phaeodactylum tricornutum* has proven to be a robust model for research, yielding a wealth of knowledge and advancing our understanding in this field of research.

*P. tricornutum*, a marine pennate diatom is commonly found in coastal waters, including tidal areas, estuaries, rock pools and shallow waters exposing the species to important fluctuations in light intensity and salinity. The diatom is a well-established model with a genome that has been fully assembled and well annotated, along with an expanding molecular toolbox using the reference strain Pt1 8.6 [6–10]. Genome wide sequencing of ten accessions of *P. tricornutum* (Pt1 to Pt10) using Illumina, identified throughout the genome diverse variations, including single nucleotide and insertion deletion polymorphisms (SNPs, INDEls) and copy number variations [11]. This study provided important insights into the genetic diversity of the isolates clustering them into four distinct clades with a conserved genetic and functional makeup. Previous studies have revealed distinguishing features among different accessions. Pt4 displayed a low non-photochemical quenching (NPQ), Pt5 demonstrated higher adhesion, Pt6 exhibited substantial lipid accumulation, Pt8, Pt3 and Pt9 have different cell morphologies and Pt3 demonstrated increased tolerance to variations in salinity, among other traits [12–14].

The 10 sequenced accessions of *P. tricornutum* were mostly collected more than a century ago and have been preserved as either lab cultures, or frozen stocks in culture collections for extended periods of time. Therefore, their genetic composition may have been affected, potentially favoring genes that are adapted to lab conditions [15–20]. Seven more recent isolates were collected from the environment and sequenced, Pt11 (HongKong, China), Pt12, Pt13 (both from Bourneuf Bay, West Atlantic, France), Pt14 (Gulf of Salerno, Italy), Pt15 (East China sea), Pt16 (Helgoland, Atlantic Ocean, North Sea, Germany) and Pt17 (Banyuls Bay, Gulf of Lion, France) (Figure 1, Table S1). Assessment of genetic diversity within natural accessions of a model diatom, such as *P. tricornutum* is critical to our understanding of fundamental questions relevant to diatom’s biology and ecology. A high genetic diversity within the *P. tricornutum* species presents substantial implications, including valuable insights into their adaptive strategies across diverse ecological niches, alongside the identification of pivotal genetic determinants governing responses to environmental factors.

**Figure 1.**
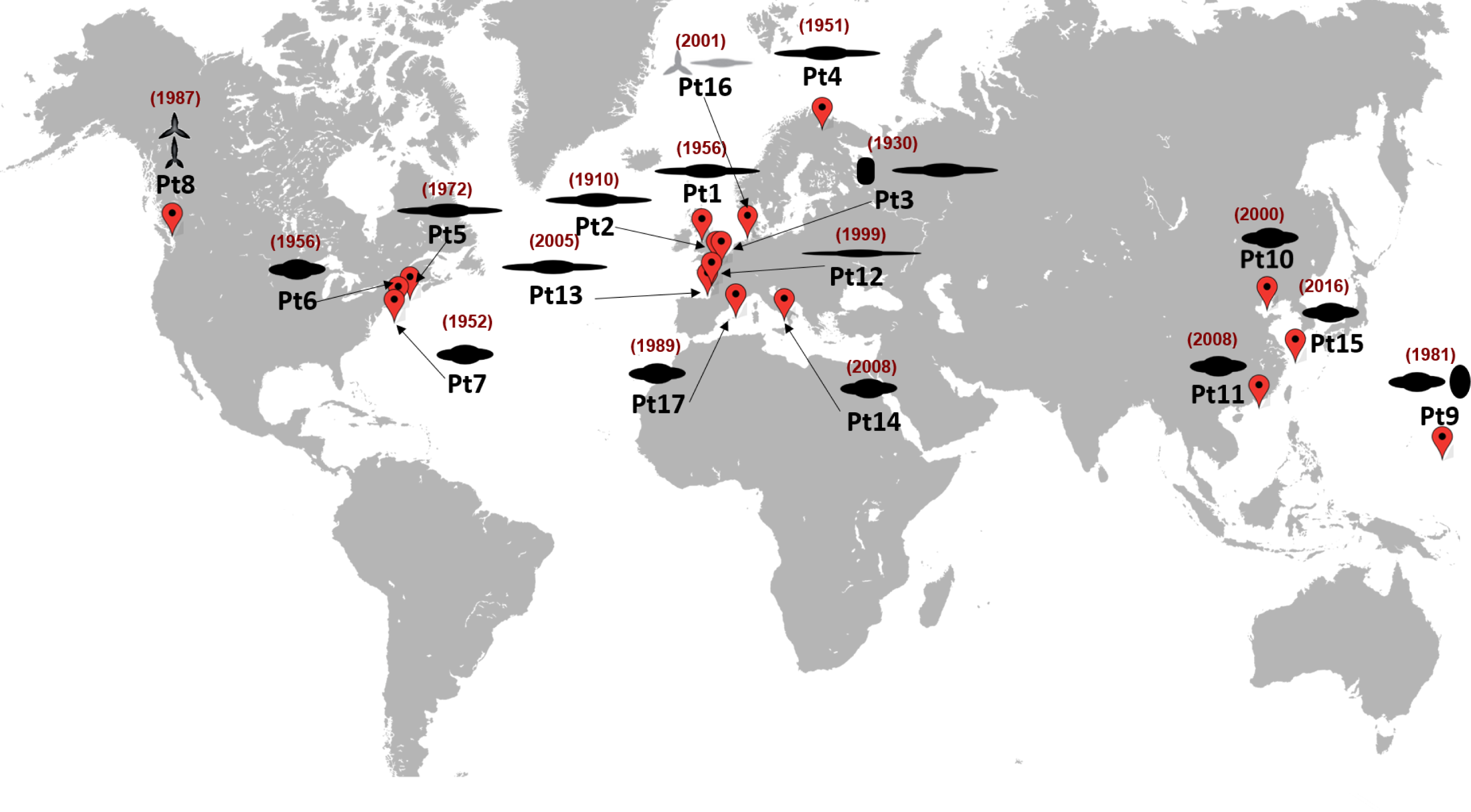
World map illustrating the sampling sites for the 17 *P. tricornutum* accessions analysed in the study with the year of sampling indicated in red.

DNA sequence polymorphism can lead to phenotypic variations but it remains only the first step in understanding how these polymorphisms can affect the phenotype. Variations in transcript levels are another proxy to better understand the contribution of genes to phenotypic variations. Diatoms have developed sophisticated sensory and gene regulatory mechanisms to detect and respond to environmental cues. They employ transcription factors and regulatory elements to control the initiation and rate of transcription. Furthermore, post-transcriptional mechanisms, such as RNA splicing, RNA transport, stability, and translation, play essential roles in determining the abundance and activity of specific gene products. These integrated processes collectively enable diatoms to fine-tune gene expression in response to changing environmental conditions [7]. An illustrative example is the diel and circadian rhythms in gene expression, which are synchronized with light and dark cycles. These rhythmic gene regulation mechanisms enable diatoms to optimize their metabolic processes and growth in a time-dependent manner, playing a significant role in their overall physiology and ecology [21].

Differences in gene expression can be attributed to different DNA sequence polymorphisms including SNPs and INDELs that can nullify gene function or induce variations in splicing. It is important to ask whether the genes that show differences in expression are under selective pressure and whether there is a link between transcript level variations and DNA sequence polymorphisms. DNA sequence may not explain differences in gene expression, cases rather known to be the consequences of epigenetic factors including DNA methylation, post-translational modifications of histones and small and long non-coding RNAs [22, 23].

In the present study, we analysed the genetic diversity of 17 accessions of the model diatom *P. tricornutum* by examining both DNA sequences and transcript levels. These accessions were collected from various coastal regions of world seas and oceans including recently sampled accessions that were not included in the previous study that had sequenced accessions collected over a span of 100 years. Our findings indicate a higher genetic diversity that defined more distinct clades and long-term balancing selection of genes in old and newly sampled accessions. However, our analysis failed to establish a clear link between the temporal factor of sample collection year and the extent of genetic diversity. Consequently, the hypothesis proposing a decline in genetic diversity, specifically the loss of heterozygosity in cultured strains over time, could not be supported [24]. Furthermore, our study identified novel transcripts among which various non-coding RNA species and provides insights into the regulation of genes mediated by genetic and transcript diversity. Our study offers easy and valuable access to these novel genetic resources, particularly focusing on a model species, through the PhaeoEpiView platform (https://PhaeoEpiView.univ-nantes.fr) [10]. This unprecedented accessibility provides a multitude of opportunities for exploring diverse ecological functions by leveraging the genetic diversity of this model organism, thereby expanding our understanding of the biology and ecology of microalgae.

## Material and methods

### Material used and growth conditions

Eighteen different accessions of *P. tricornutum* were acquired from the Provasoli-Guillard National Center for Culture of Marine Phytoplankton, Roscoff and Nantes culture collections (Table S1). All of the accessions were grown axenically using Enhanced Artificial Sea Water (EASW) [25] in batch cultures at 19°C, under 12/12 light dark period with a light intensity of 70 μmol photons m^-2^ s^-1^.

## Growth curves

The cultures were grown in 30 ml of EASW with an initial concentration of 10^5^ cells / ml. Cell counts were measured using flow cytometry (CytoFLEX, Becman, USA), every 2 days for 20 days, with 1 ml of sample taken from each accession culture each time. After 7 days of culture, 1 ml from each of Pt11 to Pt17 were used for light microscopy to describe variant shapes and cell size using Axio inverted microscope (ZEISS, Germany). Photos were further analyzed using Zeiss software (ZEN 2.6).

### Pulse amplitude modulated variable *chlorophyll a* measurements

Variable fluorescence measurements were carried out using an Imaging PAM fluorometer (Walz) with a blue measuring light (450 nm), controlled by the software ImagingWin v2.46i (Heinz Walz GmbH, Effeltrich, Germany). The actinic and saturating light were also blue and provided by fluorometer LED panel. The saturation pulse intensity was 6000 μmol photons m^−2^ s^−1^ for 0.8 s. Samples were dark-adapted for one hour before carrying out any measurements. For the construction of RLC (Rapid Light-response Curves) [26], the samples were exposed to nine incremental intensities of actinic light with an irradiance step duration of 30 s. The PAR (photosynthetically active radiation; 400 – 700 nm) steps used were: 0, 5, 19, 31, 37, 42, 47, 56, 75, 143, 280 and 519 μmol photons m^−2^ s^−1^. The first point of the RLC corresponds to the dark-adapted state, yielding the minimum fluorescence yield (*F_o_*) and the maximum fluorescence yield (*F_m_*), allowing the calculation of the maximum PSII quantum efficiency (*F_v_*/*F_m_*) as *F_v_*/*F_m_* = (*F_m_*-*F_o_*)/*F_m_*. The remaining light steps measured the fluorescence yield (*F’*), the maximum fluorescence yield (*F_m_’*) in the light-exposed state and the effective PSII quantum yield at each experimental light level (E) as *F_q_’*/*F_m_’* = (*F_m_’*-*F’*)/*F_m_’*. Relative PSII electron transport rates were calculated as rETR(E) = *F_q_’*/*F_m_’*(E) x E and non-photochemical quenching as NPQ = (*F_m_* - *F_m_’*)/*F_m_’*. Maximum relative electron transport rates (rETR_max_) were estimated by fitting the RLCs with the photosynthesis-light response model of [27] and maximum NPQ (NPQ_max_) by fitting the NPQ-light response model of [28].

### DNA extraction and PCR protocol

After 7 days of culture, cells were centrifuged at 4000 x g for 20 min and washed twice with 1XPBS. DNA was isolated using a CTAB protocol as described previously [29, 30]. A volume of 1.5 mL of CTAB buffer (450 μL 10% CTAB, 420 μL of 5 M NaCl, 60 μL of 0.5 M EDTA, 150 μL of 1 M Tris HCL) preheated to 65 °C, was placed into a 2 ml plastic tube together with diatom pellet and incubated for 1 hour at 65 °C, then DNA was isolated using Chloroform Isoamyl (24/1) after centrifugation for 10 minutes (12000 × g). The upper phase was removed and incubated with 3.2 μg of RNase A for 1 h at 37°C. DNA was isolated again using Chloroform Isoamyl (24/1) after centrifugation (12000 × g) to remove protein and RNA. Same volume of isopropanol and 8% volume of ice-cold ammonium acetate 7.5M, were used for precipitation at -20°C overnight. Nucleic acids were recovered after centrifugation (12000 × g) at 4°C and purified by absolute ethanol, then washed with 70% ethanol. DNA concentration was measured using NanoDrop ND-1000 spectrophotometer (Thermo Scientific, Wilmington, DE, USA). PCR amplification was carried out on Mastercycler^®^nexus ×2 (eppendorf, Germany) in 20 μL total volume including 9.6 μL Go Taq, 50 ng DNA and 10 μM forward and reverse primers. The PCR program consisted of 95°C for 5 min, then 35 cycles of 95°C denaturation for 30s, annealing at appropriate annealing temperature (56°C-62°C) for 30 s, 72°C extension for 30 s, and a final extension step at 72°C for 10 min. PCR products were electrophoresed on 1% agarose gel, and the gel images were acquired using EBOX CX5 System (VILBER BIO IMAGING, France). Primer sequences are listed in Table S2.

### RNA extraction

A total of 300 ml exponentially growing cells were centrifuged at 4000 x g, 4°C for 20 min and immediately re-suspended in 500 μL TRIzol^®^Reagent (Invitrogen, Thermo Fisher Scientific, USA) and vortexed vigorously before being stored at - 80 °C. RNA was isolated using Trizol reagent as described previously [31]. Purity and quantity of RNA were assessed using NanoDrop ND-1000 spectrophotometer (Thermo Scientific, USA). To remove genomic DNA, RNA samples were treated with Ambion^TM^DNase I (Invitrogen, Thermo Fisher Scientific, USA), according to manufacturer’s instructions. RNA was quantified using Qubit^TM^ RNA BR Assay Kit, 500 assays (Invitrogen, Thermo Fisher Scientific, USA).

### DNA and RNA sequencing

Extracted DNA for Pt1 8.6 and Pt11 to Pt17 was sequenced on Illumina Novaseq 6000 platform, using 250 bp paired-end reads. Yields for Pt1 8.6, Pt11, Pt12, Pt13, Pt14, Pt15, Pt16 and Pt17 were 5.9, 13.2, 6.3, 5.7, 8.0, 6.6, 5.9 and 6.6 million read pairs, respectively. Messenger RNAs for Pt1 8.6 and Pt1 to Pt17 were sequenced in duplicates on Illumina Novaseq 6000 platform, using 150bp paired-end reads. The RNA libraries were enriched for matured RNAs and sequenced in stranded mode, yielding between 15.1 and 25.7 million read pairs.

### Bioinformatics analysis Variant calling analysis

Paired-end Illumina libraries from each ecotype (Pt1 to Pt17) were first trimmed using Trimmomatic [32] with “ILLUMINACLIP:adapters.fa:2:30:10:2:keepBothReads LEADING:20 TRAILING:20 SLIDINGWINDOW:4:20 MINLEN:40”. Reads were then mapped with BWA-mem2 2.2.8 [33] on *P. tricornutum* Phatr2 assembly (accession GCA_000150955.2). Mapping rates ranged from 94.70% for Pt8 to 99.36% for Pt13. Variant calling and filtering were performed with GATK package version 4.2.2.0, following GATK best practices [34]. In short, HaplotypeCaller module was run with a call confidence of 30, a sample ploidy of 2 and double precision was activated for pair-HMM algorithm. Variants that were called were functionally annotated by snpEff [35] with *P. tricornutum* database v5.0 and transposable elements annotation from [7], an upstream/downstream region size set at 2kb and gene putative loss of function (LOF) annotation activated. Only SNPs and insertions/deletions (INDELs) were retained from annotated VCF files. Finally, GATK’s VariantFiltration module was used on SNPs with the following filters: “QD<2.0; QUAL<30; SOR>3.0; FS>60.0; MQ<40” and on INDELs with: “QD<2.0; QUAL<30; FS>200.0”. INDELs of size above 50bp were extracted with SelectVariant module and some of the longest were validated by PCR.

### Fixation index computation

Fixation index (Fst) was computed with ANGSD 0.939 [36] between all 17 accessions possible pairs. First, in order to compare only regions where data were present for all samples, the callable genome size was defined where read coverage on reference was no less than 10X across all accessions. Allele frequencies were computed for all ecotypes using ANGSD “-doSaf 1 -GL 2 -minMapQ 1 -minQ 20”, then folded site frequency spectrum (2DSFS) was determined with “realSFS -maxIter 100 -fold 1” for all ecotypes combinations. Finally, we computed all pairwise Fst on the resulting indexes with “realSFS fst index -fold 1”. All Fst values were gathered in a matrix and displayed as a heatmap using the R [37] package Pheatmap 1.0.12 (https://scicrunch.org/resolver/RRID:SCR_016418).

### Population clustering

Callable SNPs and INDELs were analyzed with ADMIXTURE 1.3.0 [38] with a random seed of 12345679, cross-validation (CV) activated and a bootstrap value of 200. Numbers of ancestral populations (K-value) were tested from 1 to 17 and cross-validation error was plotted in order to select K leading to the lowest CV error. A PCA was then performed on Q-estimates for K=15 and estimated ancestral fractions were plotted with R.

### CNV and gene loss analysis

For each ecotype (Pt1 to Pt17), raw number of mapped fragments from the Variant Calling Analysis BAM files were counted on each Phatr3 gene [7] using featureCounts [39] in unstranded paired-end mode and reads were assigned to all their overlapping features (“-O” option). Genes with no counts were deemed as possibly lost. Raw counts were then normalized for each gene following FPKM formula: FPKM_normalized_count = (gene_raw_count x 109) / (gene_length x total_sample_counts). Similar to previous work [11], binary logarithm Fold Change (log2FC) was calculated as the log2 ratio of normalized count for each gene to the average (mean) normalized count of all the genes per accession. Genes with a log2FC >= 2 were considered as showing putative Copy Number Variation (CNV) compared to the reference strain. Finally, lost genes and genes exhibiting CNV in only one accession were marked as ecotype-specific. Heatmap plots were made in R using Pheatmap 1.0.12 (https://scicrunch.org/resolver/RRID:SCR_016418) and UpsetR 1.4.0 [40] packages.

### Genes with loss of function

After variant annotation by snpEff, we used an in-house script to find the total and specific number of genes affected by loss-of-function (LoF) mutations for each ecotype (Pt1-Pt17). First, we selected genes with LoF variants alleles retained in the VCF annotation file whether they are homozygous or not. Then, we searched for the genes that are specific to each accession and considered the non-accession specific genes common if shared by two or more accessions.

### Site Frequency Spectrum (SFS) analysis

A matrix of allele counts per ecotype was created from the variants called previously on the callable genome. Briefly, for each biallelic SNP, a value of 0, 1 or 2 was determined for each ecotype, depending on its ploidy (homozygous on reference allele, heterozygous reference/alternate alleles or homozygous on alternate allele, respectively). Moreover, functional annotation as described previously and the affected gene (if applicable) were added for each SNP. Folded SFS was then calculated for each functional category of SNPs (nonsense, non-synonymous, synonymous, intergenic) and the resulting data was plotted with R.

### Searching for signatures of Balancing Selection (BS) on non-synonymous SNPs

One of the signature of balancing selection is the excess of nonsynonymous polymorphisms segregating at intermediate frequencies [41]. Genes with less than 10 synonymous (S) + non-synonymous (NS) SNPs were filtered out from the allele counts matrix (see “*Site Frequency Spectrum (SFS) analysis*”). The ratio non-synonymous versus synonymous diversity was estimated by Watterson’s theta θ assuming twice as many non-synonymous than synonymous sites (θ*w*NS/ θ*w*S), defined as “(Number of NS/2) / (Number of S)” was calculated for each of the remaining 9267 genes. The 91 genes with an excess of non-synonymous SNPs (showing a θ ratio over 3), were extracted and further investigated. Finally, the same process was performed ecotype-wise, with a number of genes with a θ ratio > 3 ranging from 34 in Pt14 to 62 in Pt4.

### Phylogeny of the 17 accessions

A matrix of genome-wide biallelic SNPs and INDELs allele counts per ecotype (0, 1 or 2 depending on the called ploidy of the variant, see “*Site Frequency Spectrum (SFS) analysis*”), was computed for 640,454 variants found in the population Pt1 to Pt17. Canberra distance and average linkage functions were identified to produce the tree that represented best the matrix by the “find_dend()” method from R library “dendextend” 1.16.0 [42]. Then, an unrooted neighbor-joining tree was built using “phangorn” R package 2.10.0 [43] on the Canberra distance matrix and colored according to the ecotypes clades.

### Expression analysis

After filtering raw data with the removal of adapters and low quality reads, clean reads were aligned against the reference genome using HISAT2 2.0.5 [44]. Reads were assigned to each transcript using the FPKM metric which normalizes for differences in library size and gene length. In order to compare gene expression levels in different accessions, the graphical representation of the distribution of gene expression and FPKM levels in different samples has been performed using the ggplot2 R package (v3.4.0)[45]. In order to differentiate between coding and non-coding transcripts, the Coding Potential Assessment Tool (CPAT) (DOI: 10.1093/nar/gkt006) which is a convenient and rapid method to categorize transcripts based on their coding scores, was used. This algorithm relies on specific models to assign coding potential scores to individual transcripts. In our research, we employed models from human, mouse, and zebrafish. Consequently, a table was generated, presenting the coding potential outcomes for each input transcript.

### Principal Component Analysis

To elucidate the relationships between distinct accessions, we conducted Principal Component Analysis (PCA) on the gene expression values (FPKM) of all the samples. Specifically, we first computed the average FPKM value for two replicates of each sample, followed by a logarithmic transformation of this average value (log2(FPKM+1)). Ultimately, we represented the samples according to their expression levels. In our analysis, we evaluated how well the samples were represented in PCA using the cos2 (square cosine) metric as a measure of quality. A cos2 value closer to one indicates a stronger representation of the variable by the two displayed components.

### Repeats detection in novel transcripts

Reads from both replicates of Pt1 to Pt17 RNAseq libraries were mapped with HISAT2 2.0.5 [44] using the default parameters and all the mapping information were combined. The reads were then assembled with StringTie 1.3.3b [46] and novel transcripts were kept. We screened these 656 novel transcripts for repeats with RepeatMasker (*RepeatMasker Open-4.0*. 2013-2015 ≤http://www.repeatmasker.org≥) Galaxy Tool wrapper version 4.0.9 (slow settings with matrices for 43% GC content), using a manually curated database of 71 reference transposable elements (TE) of *P. tricornutum*. RepeatMasker was run on the public server at https://usegalaxy.org [47].

### WGCNA network analysis

The network was obtained using Weighted Gene Correlation Network Analysis (WGCNA) R package (version 1.17) [48]. Before constructing the co-expression network, we filtered out genes having a row median less than 10 reads. The expression matrix was transformed with the vst (Variance Stabilizing Transformation) function from DESeq2 R package (version 1.32.0) [49]. The sequencing steps for the network construction for *P. tricornutum* accessions have been described previously [50].

For Network construction, the WGCNA R package [48] was used to identify network modules from 36 RNA-Seq datasets representing expression data from 18 *P. tricornutum* accessions (two replicates per accession). First, the quality of the raw counting matrix was checked. A hierarchical clustering analysis based on the “average” method allowed us to identify the Pt1R2 as an outlier, so this sample was filtered out from further analysis.

## Results

### Phenotypic traits characterization

To assess phenotypic differences among the 17 accessions, we monitored their growth, cell morphology and photosynthetic features. Significant differences in growth rate were recorded at day 4 of the exponential phase (Fig.2a). In most of the cultures, cell growth rate dropped after day 11 and entered the stationary phase. Pt8, Pt3 and Pt10 showed faster growth rate and higher final concentrations than other accessions, with 1165×10^4^, 1032×10^4^ and 960×10^4^ cells mL^-1^ respectively in the end of exponential phase (*p<0.05*). On the other hand, Pt4 and Pt9 showed slower growth rate than the others and the lowest final concentrations, with 419×10^4^ and 559×10^4^ cells mL^-1^ respectively.

Cell dimensions were measured for only 7 accessions (Pt11 to Pt17) and compared to the previously published Pt1 to Pt10 accessions [12]. Among all accessions, the previously measured Pt5 had the longest length (25-30 µm) and Pt14 cells were the shortest (10-15 µm). Pt12 were the thinnest cells with the lowest length/width ratio (9.9 ± 1.1) and Pt17 were the largest (4.6 ± 0.8) (Table S1, Fig. S1). Different morphotypes were observed in Pt16 (a mix of 75% fusiform and 25% of triradiate). As reported previously [11], we found few oval cells mixed with fusiform in Pt3 and Pt9 (7.3% and 7.6% respectively). Triradiate were reported in Pt8 [12] but we did not observe triradiate cells in our conditions. Triradiate morphotype was reported to be instable in this accession [51].

### Measurements of photosynthetic parameters

To assess photosynthetic capacities of *P. tricornutum* accessions, we measured maximal PSII quantum yield (*F_v_*/*F_m_*) and maximal relative electron transport rates (rETRmax). Among all the accessions, Pt12 showed the highest *F_v_*/*F_m_* followed by Pt13 (p = 0.027) and Pt11 (p = 0.0047) (Fig. 2b). These three accessions also showed the best photosynthetic performances based on rETRmax (Fig. 2c), while Pt4, Pt5, and Pt6 demonstrated the lowest values in PSII quantum efficiency (Fig. 2c). The English Channel accessions, Pt1, Pt2 and Pt3 showed similar results.

**Figure 2.**
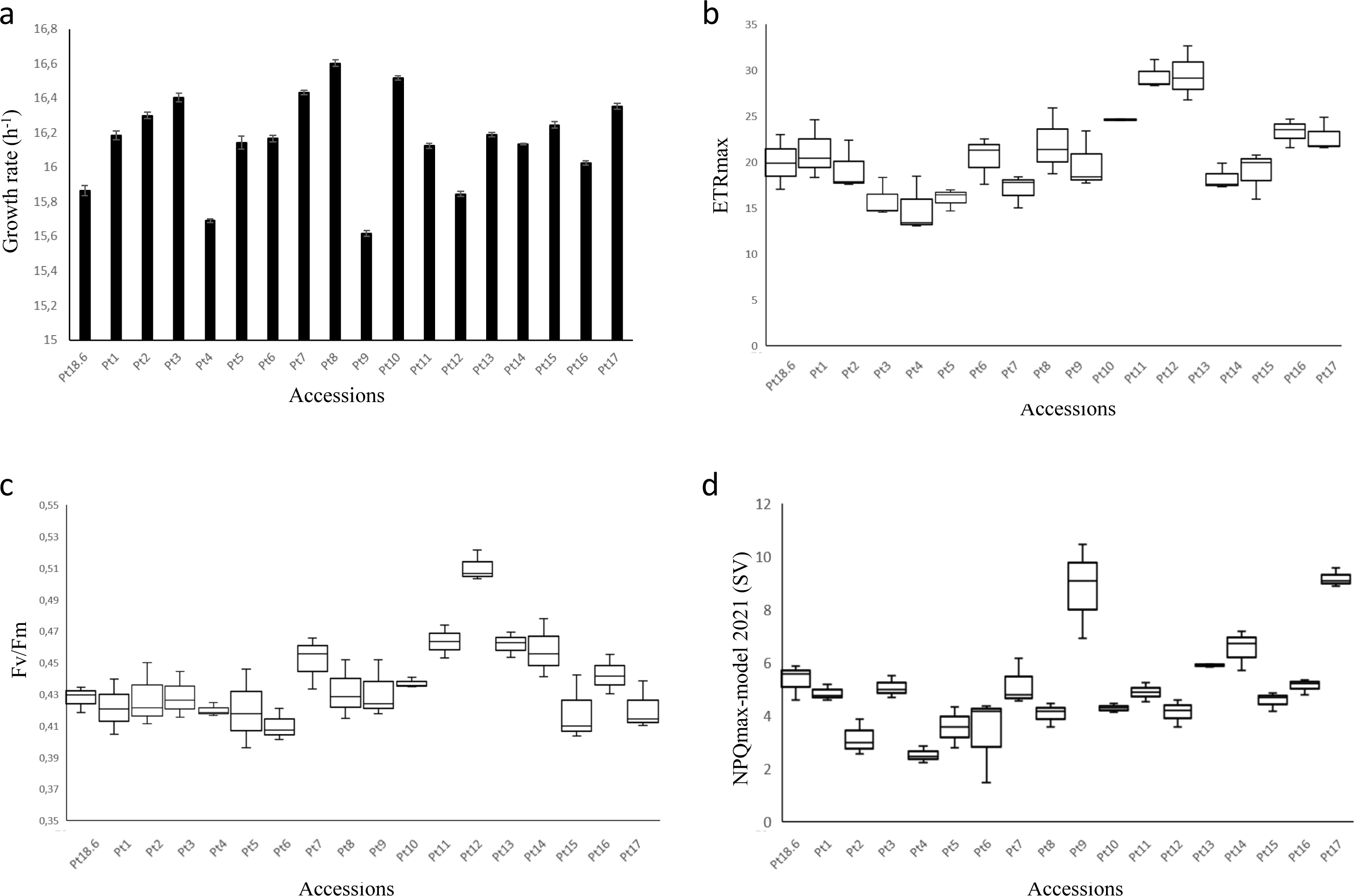
Growth and photosynthetic features for the 17 accessions. **a** Growth rates in the 17 accessions, with error bars indicating standard deviations based on triplicate cultures. **b**. Mean maximum relative electron transport rate (rETRmax) for the 17 accessions. **c** The maximum PSII photochemical efficiency (Fv/Fm). **d** Non-photochemical quenching (NPQ) for the same 17 accessions.

To evaluate the response to excess light energy, we measured non-photochemical quenching (NPQ). Pt9 and Pt17 showed the highest NPQ capacity, c. 8.2 and c. 9.2 respectively while Pt4 showed a lower NPQ capacity of around 2.5 (Fig. 2d). These observations suggest that Pt9 and Pt17 can tolerate environments with higher light intensity compared to other accessions. This is consistent with their geographical distribution in latitudes that receive greater amounts of solar radiation. Similarly, Pt4 NPQ reflects an adaptation to its sampling location in higher latitudes, specifically the Norwegian Fjords.

### Variant calling analysis

We performed variant calling analysis on previously published sequences of Pt1 to Pt10 [11] and the newly sequenced Pt11 to Pt17 accessions using the reference strain Pt1 8.6 genome sequences. All the accessions had a good sequence coverage allowing a confident variant calling. We identified 731,357 single nucleotide polymorphisms (SNPs), 44,470 insertions (from 1 to 422 bp length) and 52,867 deletions (varying from 1 to 274 bp) (Fig. 3 a,b). Site frequency spectrum (SFS) which reflects the numbers of variants segregating at different frequencies showed the expected excess of low frequency alleles (Fig. 3c). The highest increase in low frequency SNPs was observed in non-sense polymorphisms. Non-synonymous polymorphisms showed the second highest increase, when compared to intergenic and synonymous polymorphisms. Most of the SNPs (59-63%) were found in genes while INDELS were mostly detected in intergenic regions (Fig.3 d). Our analysis identified 22,4253 additional SNPs and 74,918 additional INDELs in Pt1 to Pt10 compared to our previous study [11].

**Figure 3.**
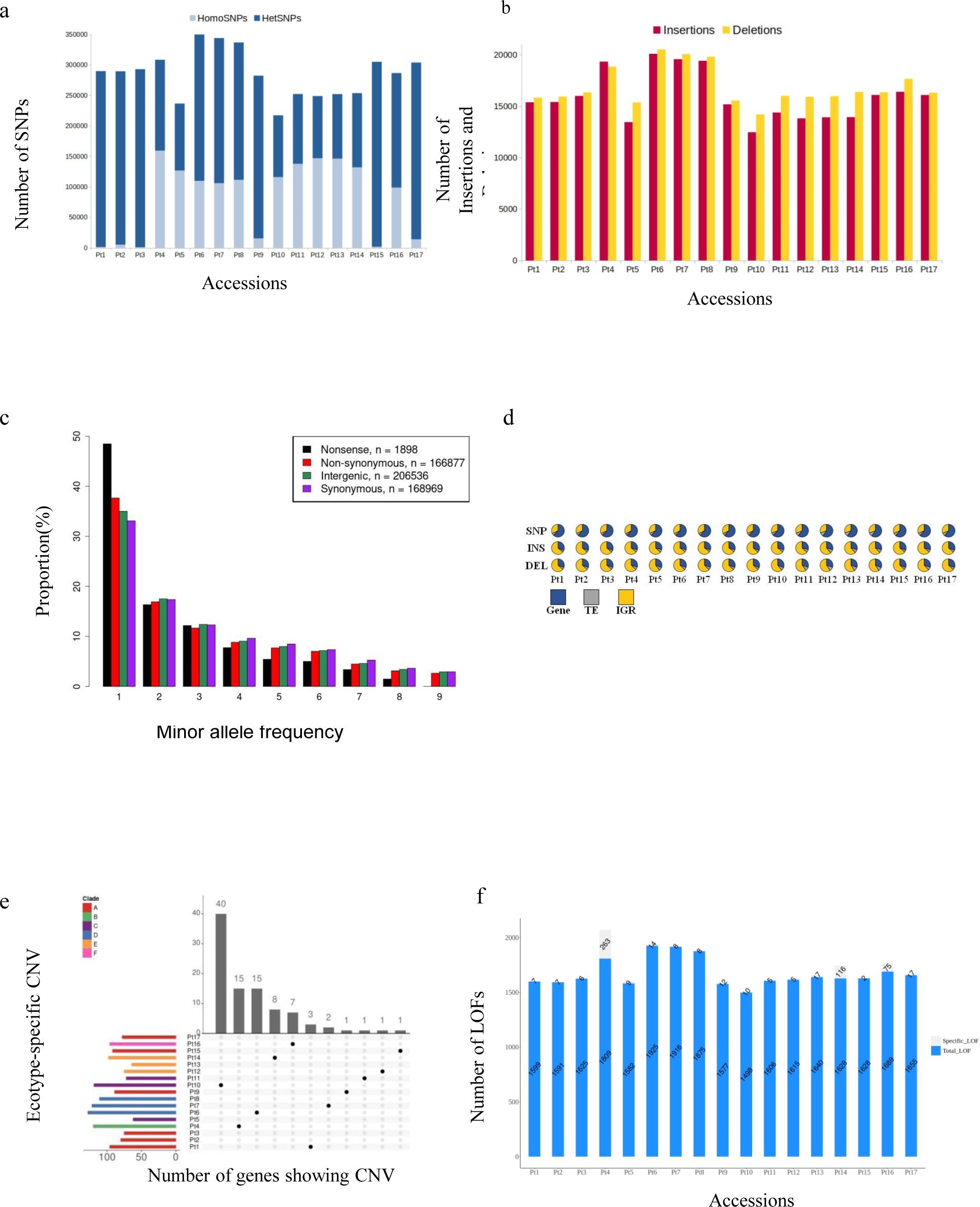
Genetic diversity across *P. tricornutum* accessions. **a** Composition of heterozygous and homozygous SNPs, the stacked bar plot represents the number of SNPs discovered in Pt1 to Pt17 accessions, showing a contrasted proportion of homozyguous and heterozyguous SNPs. Homozyguous SNPs are displayed in light blue, heterozyguous SNPs are shown in dark blue. **b** The bar plot represents the number of insertions (red) and deletions (yellow) called in Pt1 to Pt17 accessions. **c** Folded site frequency spectrum SFS of 1898, 166877, 206536 and 168969 non-sense, non-synonymous, intergenic and synonymous SNPs, respectively. In order to obtain an unbiased SFS, only one accession was chosen to represent each group of genetically close accessions. These variants were thus called on Pt1-4-7-9-11-12-14-16-17. **d** Pie charts represent different proportion of SNPs and INDELs over all functional features of the genome; GENEs (blue), TEs (gray), IGRs (Intergenic Regions, represented in yellow). **e** The bar plots represent the total number of genes considered to exhibit CNV per accession (left, colored by clade) and those that are accession-specific (top, in grey). Only accessions having specific genes with CNV are shown in the matrix (center, in black). **f** The bar chart represents total and specific numbers of genes that are affected by loss-of-function (LoF) mutations for all ecotypes (Pt1-Pt17). The total number of genes (blue color) is the number of all non-duplicated genes on which a single variant (INDEL or SNP) was taken into account. The grey bars represent the number of genes unique to a specific accession and not present in the others.

Despite the higher number of discovered SNPs and INDELs, the overall trend of their distribution among the 10 previously analyzed accessions remains the same. Across all the accessions most of the SNPs were heterozygous (HetSNPs) with >95% in Pt1, Pt2, Pt3, Pt9, Pt15 and Pt17 and 65% to 68% in Pt6, Pt7, Pt8 and Pt16 and the lowest proportion of HetSNPs were found in Pt4, Pt5 and Pt10 to Pt14 (<49%). Across all the accessions, the numbers of INDELs was similar except for Pt4, Pt6, Pt7 and Pt8, which showed the highest number of INDELs (Fig. 3b, Table S3). Most INDELs were shared among the accessions. A total of 14 INDELs were validated by PCR for randomly chosen loci (Fig. S2). To further assess genetic diversity, we investigated copy number variations (CNVs) and gene loss (GL). A total of 284 and 180 genes show CNV or GL respectively. Most of CNVs are shared and eleven accessions out of 17 show specific CNVs with 40 genes in Pt10 followed by Pt4, Pt6, Pt14 and Pt16 with 15, 15, 8 and 7 genes respectively (Fig. 3e, Table S4, Table S5). Randomly chosen loci were validated by PCR for gene loss (Fig. S3).

To understand the functional impact of genetic variations among *P. tricornutum* accessions, we examined the loss of function mutations (LoF) such as premature stop codons, frameshifts and start loss. A total of 31,536 LoF was found, among which 588 were shared across the accessions (Fig. 3f). Accession specific LoFs were mostly found in Pt4 (61), Pt14 (20) and Pt16 (17) (Table S6). LoF mutations were enriched in gene ontology (GO) categories of molecular function type (Table S6) and in genes that belong to large gene families as previously shown [11].

### Population structure and phylogeny of *P. tricornutum* accessions

To examine the global population structure of *P. tricornutum* accessions, we used pairwise fixation index (Fst), a measure of genetic differentiation revealing groups with low Fst index (< 5%) (Figure 4a). To further estimate genetic relatedness among *P. tricornutum* accessions, we used admixture proportion inference which allows the assignation of individual genetic variations into clusters based on shared allele frequency patterns [38]. We ran ADMIXTURE with various plausible values of *K* which is the number of source populations and found a stable admixture pattern proportions with K=15, reflecting the number of ancestral populations (Fig. S4, Table S7). Based on individual ancestry with similarity of clusters between accessions, we distinguished 6 clades: Pt1, 2, 3, 9, 15 and 17 in clade 1 with most of the clusters (up to 11) reflecting a clade with multiple ancestral populations, Pt16 in clade 2, Pt4 in clade 3, Pt5, 10 and 11 in clade 4 with only 3 clusters, Pt6, 7 and 8 in clade 5 and Pt12, Pt13 and Pt14 in clade 6 (Figure 4b).

**Figure 4.**
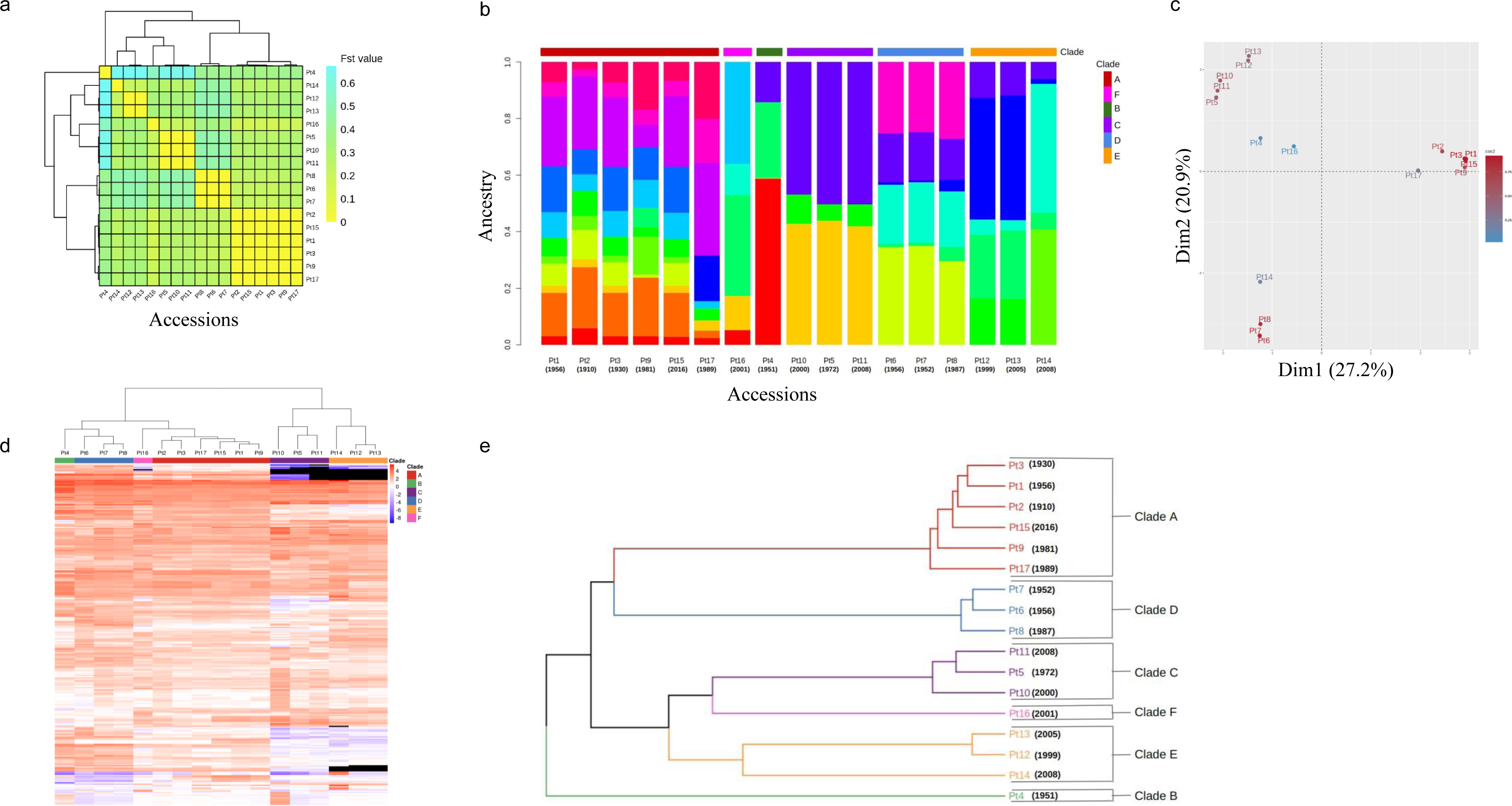
Clustering of *P. tricornutum* accessions into clades. **a** The heat-map shows the genetic differentiation or association between all possible pairs of accessions. The colors indicate Fst values, which range from 0.02 to 0.4, with a color gradient from yellow to green, respectively. Values closer to 0 signify close genetic makeup and values closer to one indicate strong genetic structuring between the populations. **b** ADMIXTURE plot representing the ancestry genome fractions of Pt1 to Pt17 (in color) for K=15. **c** Principal component analysis (PCA) showing the distancebetween the seventeen accessions based on their shared genome structure. **d** The heatmap shows the log2 fold change (log2FC) of normalized reads counts between each reference gene and average of the read counts of all the reference genes per accession. FPKMs are used as normalized values, the log2 ratio of each gene FPKM over the mean FPKM of each accession being calculated. A blue to red color gradient in the heatmap represents low to high log2FC. The previously described clades A, B, C, D and the newly defined clades E and F are shown as colored annotations on the top. Genes having a null FPKM in a given accession (considered lost) are displayed in black. Only genes having a log2FC over 2 in at least one ecotype are plotted in this figure (222 genes) and are considered to exhibit Copy Number Variation (CNV). **e** Phylogenetic association of the 17 accessions based on genome-wide biallelic SNPs and Indels (640454 variants), built from a hierarchical clustering (canberra distance and average linkage functions).

To confirm admixture analysis clades, we performed a Principle Component Analysis (PCA) which revealed similar results reflecting common ancestry except for Pt14 which is far from its admixture defined clade 6 (Fig. 4c). Of note, cluster composition proportions of Pt14 are different from Pt12 and Pt13. To further assess the segmentation among *P. tricornutum* accessions, we used CNVs to run a hierarchical clustering and found that the 17 accessions fall into 6 clusters supporting further both PCA and admixture analyses: Pt1, Pt2, Pt3, Pt9, Pt15 and Pt17 in cluster 1, Pt16 in cluster 2, Pt4 in cluster 3, Pt6, Pt7 and Pt8 in cluster 4, Pt5, Pt10 and Pt11 in cluster 5 and Pt12, Pt13 and Pt14 in cluster 6 (Fig. 4d). Phylogenetic analysis at whole genome scale using genetic variations (SNPs and INDELs) of the 17 accessions supported further the clustering into six clades observed with Fst and PCA analyses (Fig. 4e).

### Balancing and relaxed selection in Pt clades

We calculated the ratio of nonsynonymous to synonymous nucleotide site diversity using Watterson’s estimate of theta (θ*w*N/θ*w*S) [52] as a measure of the efficiency of natural selection. Ratios > 3 suggest that selective pressure maintains non-synonymous polymorphisms, a signature for balancing selection (BS) which refers to selective processes by which alleles are maintained in a population at frequencies larger than expected from genetic drift alone [53], while θ*w*N/θ*w*S ratios < 1 refer to non-synonymous polymorphisms being counter-selected pointing to a purifying selection. We identified 91 common genes under BS and 2422 under purifying selection (Table S8). Most of the genes under BS are of unknown function. However, those with known function were enriched in genes coding for functions such as cell proliferation and growth, perception, transmembrane transport activity, stress responses and adaptation to the environment.

### Identification of transcript level variations and co-expression network modules in *P. tricornutum* accessions

Differences in gene expression is known to control inter and intra specific phenotypic variations providing living organisms with abilities to colonize different ecological niches. To identify differences in transcriptomes, RNAs from *P. tricornutum* accessions (Pt1 to Pt17) were sequenced and mapped to the reference genome. The mapping of mRNA-Seq reads was above 85% for all replicates except for one, Pt16R2, which had a mapping rate of 41.27%. Pearson correlation coefficients between each of the two replicates was mostly around 0.98 (Fig. S5). Interestingly, a total of 656 genes including 25 from the chloroplast were novel. These genes are widely distributed over the genome among which some were specific to each of the ecotypes and 385 genes were found to be common to all of them (Table S9). They showed an average length of 709.42 bp, with334 genes categorized as sense and 322 genes as antisense. Except from few genes that were annotated, most of these novel genes were of unknown function (Table S9). The majority of novel genes were downregulated compared to the average gene expression but their expression remains significant and cannot be considered as part of a spurious phenomenon of background low-level transcription.

The analysis of novel transcripts using RepeatMasker revealed that 12.26% of them contained repetitive elements, primarily Copia LTR_retrotransposon of class I and rare MuDR2 Terminal Inverted Repeats of class II (Table S10). Additionally, approximately 0.48% of the transcripts contained simple repeats. Using a coding potential assessment tool, we identified several non-coding RNA with sizes varying between 202 bp and 8379 bp (Table S11). Furthermore, we detected several other RNA types, including sRNA, miRNA, tRNA, snoRNA, antisense and piwiRNAs (Table S11).

Then, we examined differentially expressed genes (DEGs) under our standard growth conditions by analyzing RNA-Seq data across the 17 accessions and comparing them with the reference strain Pt18.6. This strain was derived from Pt1, which displayed the lowest number of DEG (1308 genes) as expected (Figure 5 a). In contrast, Pt7 showed the highest number of DEG (5086 genes). The remaining ecotypes showed variable numbers of DEG, with Pt5 exhibiting the lowest number at 1890 genes. On average, most ecotypes had approximately 4000 DEG. With the exception of Pt1, which exhibited a bias towards upregulation (twice as many upregulated genes as downregulated genes), the other ecotypes displayed a balanced ratio of upregulated and downregulated genes. The majority of upregulated genes, their log2(FoldChange) values varies between 1 and 3,while the majority of downregulated genes,the value of log2(FoldChange) varies between -3 and -1. A substantial fraction of DEGs, regardless of their upregulation or downregulation status are found to be specific to one or multiple ecotypes suggesting an ecotype-dependent gene expression pattern that likely underlies ecotype specific trait (Figure 5b, c, Table S12, S13). On the other hand, only a minor subset of DEGs displaying upregulation or downregulation were shared across all the ecotypes. We conducted a principal component analysis using the average replicates expression to evaluate whether the ecotypes exhibited comparable expression profiles. Our analysis distinguished five clusters that partially aligned with the clades defined in this study, implying a correlation to some extent between genetic diversity and expression patterns (Figure 5d).

**Figure 5.**
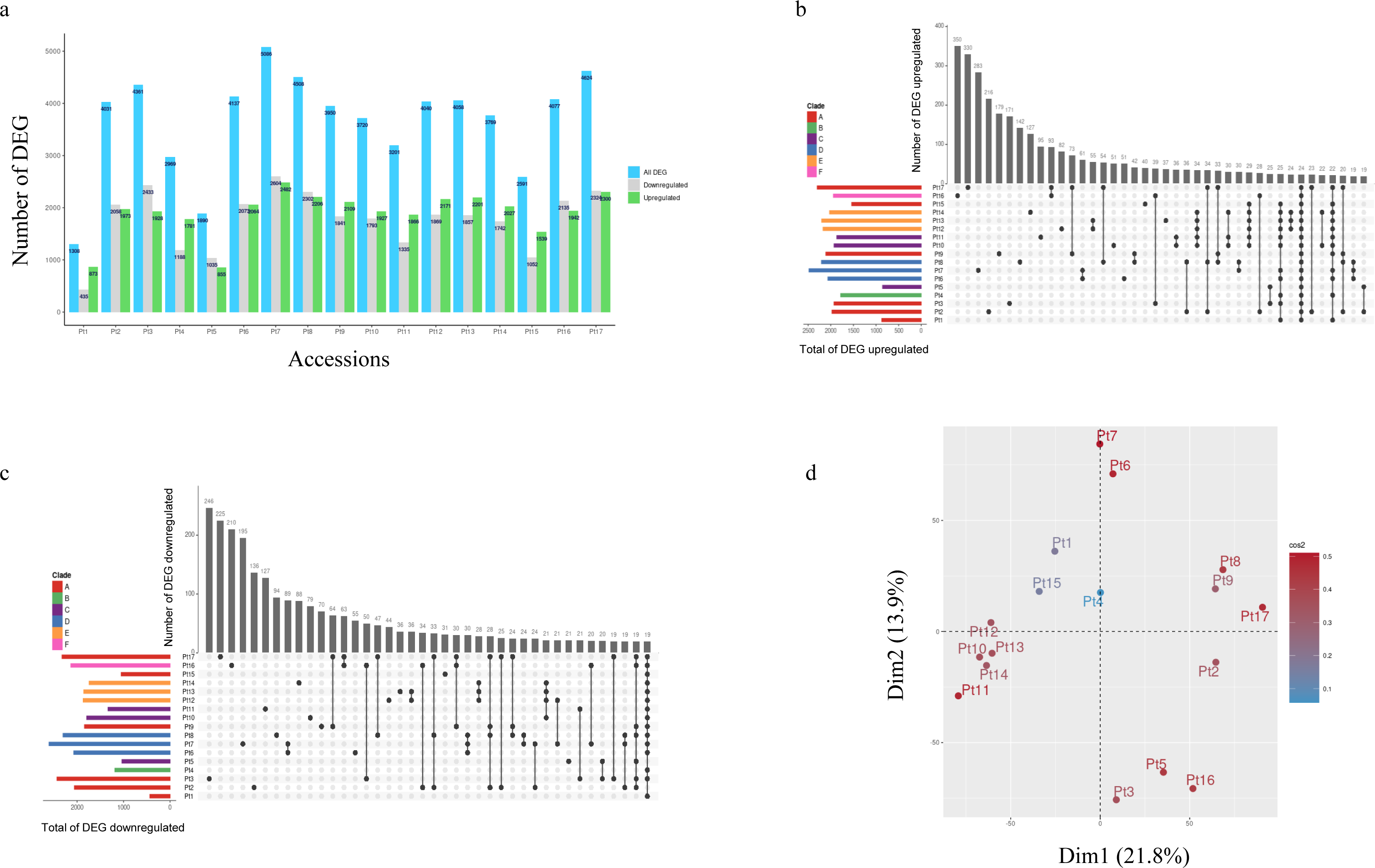
Transcription levels variations in the 17 accessions. **a** The bar plots represent the number of differentially expressed genes (DEG) in the 17 accessions denoted on the Y axis. All DEGs are displayed in blue. Downregulated genes are shown in grey and upregulated in green. **b** The bar plots represent total number of upregulated genes in each accession (left, colored by clade) and those that are specific to a given group of accessions (top, in grey). **c** The bar plots represent the total number of downregulated genes in each accession (left, colored by clade) and those that are specific to a given group of accessions (top, in grey). **d** Principal component analysis (PCA) showing the distancebetween the seventeen accessions based on gene expression values.

To explore the relationship between phenotypic traits, specifically photosynthesis and transcripts, we closely examined the NPQ response and the genes related to its regulation in response to light. Indeed, the NPQ capacity depends on transthylakoidal proton gradient, but also on antenna proteins called Light-Harvesting Complex Protein X (LHCX) and on the Diatoxanthin Xanthophyll cycle [54], equivalent to the Zeaxanthin cycle in land plants [55]. Among the LHCX genes, only LCHX1 shows strong expression in all ecotypes, confirming its constitutive role in low light, while LHCX2 and 3 are involved in high light and LHCX4 expression increases in the dark [56] (Fig. S6). Similarly, all genes involved in the Diatoxanthin Xanthophyll cycle show negligible expression in our low light culture conditions.

To understand the underlying nature of the conserved transcriptomic responses, we analyzed the enrichment of GO terms for both upregulated and downregulated DEGs (Figure S7). Additionally, we performed GO enrichment analysis on genes that were specifically upregulated or downregulated in a single accession as well as per clade, where applicable. Only few GO terms emerged from the analysis of accession specific DEGs, namely photosynthesis GO related terms (light harvesting, protein chromophore linkage) in Pt8, lipid metabolic processes and translation in Pt3 for downregulated genes, while upregulated genes were enriched in ribosome biogenesis and rRNA processing in Pt7, protein transport in Pt16 and glucose metabolism in Pt11 (Figure S7, Table S12, S13).

The WGCNA package was used to construct gene co-expression network of transcripts from an expression matrix of ∼432,000 transcripts derived from 36 RNA-seq samples, with 2 replicates collected from the 18 accessions including the reference strain Pt1 8.6. This approach yielded in 33 distinct co-expressed modules (labeled by different colors) with dark slate blue and plum2 modules containing each the smallest number of genes (106) and green yellow with the largest number of genes (1599) (Figure 6c, Table S14). These modules were constituted by genes demonstrating analogous expression profiles, which may or may not be consistent among different clades suggesting the existence of additional factors besides genetic polymorphisms that could modulate transcript levels (Figure S8). The modules were further categorized into six distinct clusters, each characterized by a group of genes exhibiting comparable expression patterns thus implying their involvement in shared pathways (Fig. 6c, Table S15). Based on the GO annotation analysis, we identified significant functional enrichments in different groups. Group I displayed a substantial increase in oxidoreductase activity, while Group II showed an enrichment in calcium ion binding activity. Group III was characterized by an enrichment in chlorophyll binding and light harvesting, whereas Group IV was associated with RNA processing and translation. Group V showed a significant enrichment in photosynthesis and cell redox homeostasis, while Group VI exhibited an enrichment in cell division and DNA binding activity (Table S15).

**Figure 6.**
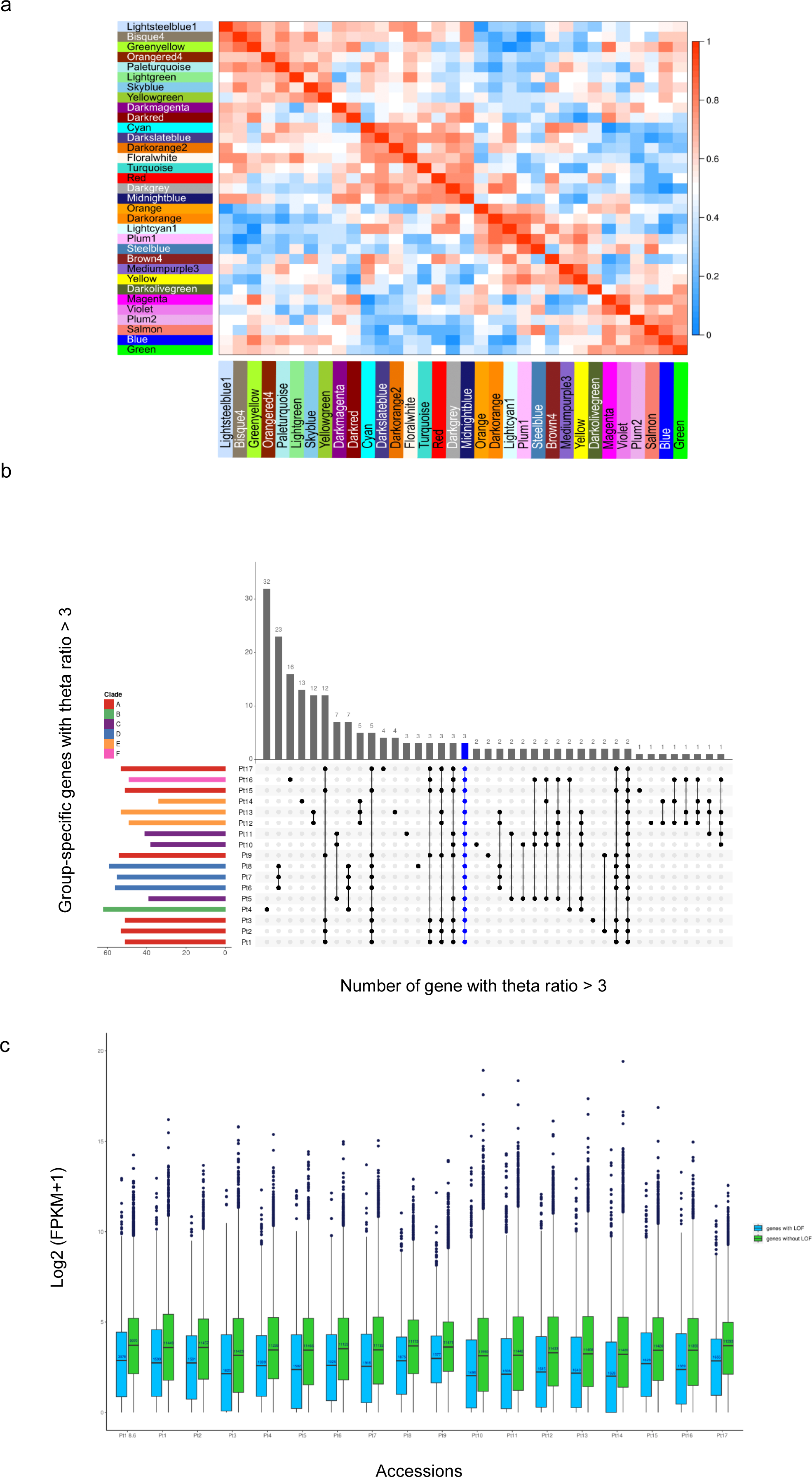
Balancing selection, LOF and WGCNA analyses. **a** The plot represents the total number of genes showing a signature for balancing selection (BS) per accessions (left, colored by clade) and those that are found only in a given group of accessions (top, in grey). Genes under BS in all accessions are shown in blue. **b** Distribution of gene expression levels in genes affected by loss-of-function mutations (blue) and unaffected genes (green) for each accession using FPKM values. **c** Eigengene adjacency heatmap of the 33 merged modules of *P. tricornutum* accessions network. Each row and column in the heatmap corresponds to one module (represented by their color). The scale bar on the right represents the correlation strength ranging from 0 (blue) to 1 (red).

## Discussion

The present study was designed to comprehensively characterize the phenotypic, genetic, and transcriptomic diversity among seventeen distinct *P. tricornutum* accessions, which were collected from various locations across the world’s oceans and included more recently sampled accessions compared to those examined in prior studies [11, 12]. Growth dynamics, cell morphology, and photosynthetic traits were monitored, and significant inter-accession differences were recorded. Notably, the Pt4 and Pt9 strains exhibited a distinct growth pattern, implying a probable correlation to their respective sampling locations. More specifically, Pt4, sampled from the Norwegian fjords, seems to be adapted to a lower light intensity regime than that employed in our study, while Pt9, a tropical strain, showed a slower growth at 19°C compared to the presumed higher temperature in its geographical location. Moreover, the evaluation of photosynthetic abilities across various accessions corroborated the association with the sampling sites. Pt4 displayed the smallest rETR_max_ and PSII maximum quantum efficiency, while Pt12, Pt13, and Pt14 collected from the Mediterranean Sea and Atlantic side demonstrated higher photosynthetic performance, as evidenced by their important F_v_/F_m_ and rETR*_max_* values. Strains within the same clade show similar growth and photosynthetic performances, but there is no apparent correlation pattern with the year of sampling. Each clade includes accessions from both older and more recently collected samples. Measuring cell dimensions in the recently acquired accessions and their comparison with previously characterized ones revealed notable variations. Specifically, Pt12, Pt14, and Pt17 exhibited significant deviations, with Pt12 having the shortest cell length, Pt14 displaying the smallest length-to-width ratio, and Pt17 showing the largest cells when compared to the remaining accessions. Additionally, our investigation identified Pt16 as a new accession with a combination of triradiate and fusiform morphotypes. These disparities in cell sizes may potentially confer an advantageous trait in terms of enhanced gliding capabilities and improved photosynthetic efficiency [57]. The observed differences in cell dimensions and, at times, morphology are not surprising, given that *P. tricornutum* does not rely on silica for growth. This lack of dependence on silica may confer flexibility in morphogenesis, a trait not typically observed in silicified diatom species. The variations in cell sizes signify an adaptation to local environments, highlighting an environmental-induced control of morphogenesis rather than a genetic one, as demonstrated previously [51]. Drill-core records from Lake Titicaca in Peru revealed a strong correlation between size trends in the diatom *Cyclostephanos andinus* and environmental variables. This suggests that diatom size responds to regional environmental changes driven by global processes that affect lake level and thermal stratification [58]. This, in turn, implies that environmentally mediated epigenetic changes modulate phenotypic traits within the same species

Variants calling analysis showed a larger number of SNPs and INDELs than previously reported in Pt1 to Pt10. This is due to the gapped alignment mode used for SNPs and INDELs calling which performs better than ungapped mapping [59]. Improvements made to HaplotypeCaller’s algorithm since 2018 were also likely playing a role in the gain of sensitivity we noticed. Most of the SNPs were located in coding regions, while INDELs were mostly found in intergenic regions as a consequence of their highly deleterious effects within coding regions. Most of the SNPs were found to be heterozygous, indicating that *P. tricornutum* has a high level of heterozygosity. Our previous study has demonstrated a substantial level of heterozygosity in *P. tricornutum* populations. The recent sampling of genetically distinct accessions has reaffirmed the persistence of this trait, despite not being related to the previously identified highly heterozygous accessions Pt1, Pt2, and Pt3, thereby supporting the reliability of the heterozygosity measure. This high level of heterozygosity is intriguing in *P. triconrutum* considering that the species is not known to reproduce sexually suggesting an advantage of heterozygous alleles or the detrimental homozygous alleles that get selected against, as reported in inbred population of clonal honey bees, *Apis mellifera capensis* which retained after 20 years of inbreeding high heterozygosity throughout its genome due to selection against homozygotes [60]. Similar examples of heterozygosity advantage through its maintenance at high proportions of the genome were reported in other species, isolated wolf populations and a hermaphrodite worm after several generations of selfing [61, 62]. An alternative explanation for the observed high heterozygosity could be due to the mutations that occurred in the ancestral lineage of Pt1, Pt2, Pt3, Pt9, Pt15, which was revealed through admixture analysis, indicating that these accessions share a common ancestry and are closely related to Pt17, which also exhibits high heterozygosity. In contrast, accessions with lower heterozygosity display a distinct ancestry pattern.

The SFS analysis provided compelling evidence consistent with the predictions of the nearly neutral theory of evolution, revealing an excess of low frequency alleles. This prevalence of lower frequency alleles in non-sense and non-synonymous polymorphisms can be attributed to the effects of purifying selection, which acts against deleterious mutations. Consequently, our investigation aimed to examine whether there was an elevated occurrence of non-synonymous mutations in the Pt1 8.6 reference strain, in contrast to both the original Pt1 accession and the closely related Pt2 strain within the same clade. This analysis sought to determine if the Pt1 8.6 strain had undergone a process of “domestication.” However, unexpectedly, we did not observe differences in non-synonymous mutations. Instead, we observed a remarkable predominance of LOF mutations in the reference strain Pt1 8.6, suggesting an adaptation to laboratory culture conditions facilitated through LOF mediated mechanisms. The majority of these LOF mutations resulted in the repression or reduced expression of targeted genes. However, a substantial number of genes showed moderate to high expression levels, implying that these LOF mutations may function as an evolutionary mechanism for generating new functional genes, serving as an adaptive response to culture conditions [63–65]. An illustrative example is the domestication of maize where most of the mutations are loss of function and the selection for a variety of traits has led to fixation of loss of function alleles in today’s crops [66, 67]. Another example is the human loss of function mutations in the promoter of a red blood cell chemoreceptor, DARC that resulted in the protection of human against malaria caused by *Plasmodium vivax* [68]. It is important to note that not all LOF mutations lead to complete functional knockout. For instance, LOF mutations at the 5’ or 3’ regions of genes may not entirely abolish their functions, and truncated proteins resulting from such mutations could act as dominant-negative factors [69, 70]. Interestingly, about 10.75% (331 out of 3078 genes) LOF showed moderate to high expression. Some specific examples of these genes with known functions include: (i) a mitochondrial enzyme known as glutamate dehydrogenase (Phatr3 J13951), which has been reported to play a crucial role in carbon and nitrogen metabolism as well as energy supply under abiotic stresses in *Arabidopsis* and the red alga *Pyropia haitanensis* [71–74] ; (ii) an LCH15 protein (Phatr3 J48882), which functions as a chlorophyll binding protein and potentially contributes to the adaptation to different light conditions in laboratory cultures and (iii) a heat shock transcription factor (Phatr3 J49594) which similarly may contribute to the adaptation to lab culture temperatures.

Admixture analysis, PCA, and hierarchical clustering all identified six clusters, which suggests that the samples had shared ancestry and similar geographical origins. However, not all accessions within the same cluster had shared geographical origins. The English Channel and East China Sea populations clustered together, indicating that *P. tricornutum* accessions may have been dispersed by various means, or that similar ecological niches across the sampling sites led to convergent evolution. Genome-wide phylogeny analysis confirmed six clades, with some accessions falling into previously identified clades and others forming two new clades, which suggests genetic divergence.

Several loci that are believed to be under balancing selection were found to have a high level of genetic diversity. Notably, genes coding for stress-inducible proteins (Phatr3_EG00471, Phatr3_J54019), outer membrane receptors (Phatr3_EG01193), and cell cycle genes (Phatr3_J34920, Phatr3_EG00817) displayed an excess of polymorphism, reflecting their role in protecting cells from stresses such as high temperatures, starvation, and infection, as well as in cell division and growth. Consistent with this, transmembrane protein have been previously observed to evolve faster than protein without a transmembrane domain in unicellular eukaryotes such as yeast [75] and *Ostreococcus* [76]. Interestingly, the genes identified under balancing selection were found in multiple clades and were specific to one or more accessions, suggesting that spatially varying selection forces may be related to local niches. It is expected that these same selection forces will apply to accessions from similar ecological niches or with a shared origin and sampling locations.

Comparison of genes under BS between accessions sampled at divergent time points, namely 1910, 1930, 1956, 1989 and 2016 revealed several identical genes suggesting the persistence of long term balancing selection acting on these genomic regions (Figure 6a, Table S8). Notably, the majority of these genes were functionally associated with stress resistance and fundamental cellular processes, highlighting their potential significance in adaptation to various habitats. Long term balancing selection was found at genes involved in diverse processes such as disease resistance, self-incompatibility, and heat stress providing advantages and enhancing fitness in natural populations [77–80].

Profiling transcript levels in the 17 accessions identified novel genes in the assayed growth conditions suggesting condition and accession specific genes that were not identified in the numerous growth conditions previously reported [7]. Interestingly, our analysis revealed that genes carrying LOF mutations displayed a significant decrease in expression levels when compared to their non-LOF counterparts (Figure 6b), implying the role of DNA sequence variations in shaping gene expression patterns. Nonetheless, we also noticed a considerable number of LOF mutations that did not result in downregulation. The observed LOF mutations with no effect on gene expression is likely due to the robustness of the genome through gene duplication and compensatory mechanisms allowing for the tolerance of many LOF variants, resulting in the majority of these mutations being silent and having little to no impact. Multiple LOF mutations were observed across clades as well as within them, particularly among accessions that exhibited extreme phenotypes (short cell size versus long ones, low versus high photosynthetic performance). This suggests that there may be variations in genetic backgrounds and/or epigenetic factors among these accessions. For instance, Pt12 and Pt14, despite having vastly different cell morphologies (very long versus short cells), share 1273 LOF mutations. Similarly, Pt2 and Pt9, as well as Pt6 and Pt12, share 1479 and 1120 LOF mutations respectively, but exhibit distinct photosynthetic performances. In general, no clear association can be established between genetic diversity including SNP, INDELs and LOF mutations and the phenotypic traits investigated in this study.

Interestingly, we observed LOF mutations that led to upregulation of genes instead, suggesting the potential existence of Gain of Function (GOF) mutations. These GOF mutations known to occur mostly in unstructured regions may be attributed to the emergence of transcription factor binding sites, miRNA binding sites, an RNA binding protein or new functional domains [81, 82]. Since genes and their products do not operate in isolation but rather in biological networks, these newly acquired domains may acquire functionality through their interactions with other proteins. Additionally, compensatory mechanisms for LOF or GOF may involve epigenetic processes that serve as a platform for interacting with specific proteins or complexes. WGCNA analysis revealed several network modules that were further merged into six clusters with similar expression patterns indicating co-regulated genes and pathways. Both differential expression and WGCNA analysis corroborated the presence of differences in transcript levels, which cannot be solely attributed to genetic variations. This observation implies the involvement of other regulatory mechanisms, such as epigenetics, that are known to modulate gene transcription [6, 83, 84].

### Conclusions

Our study provides a comprehensive assessment of the genetic and transcriptional diversity among 17 natural accessions of the model diatom *P. tricornutum*. Our investigation has uncovered novel clades, which are likely indicative of previously unexplored ecological niches. Moreover, we have identified new genes that expand the existing transcriptome repertoire of this species. By incorporating recently sampled accessions, we have further confirmed a persistent long-term balancing selection and the high level of heterozygosity in *P. tricornutum* through population genetic analyses, suggesting that this characteristic arises from a heterozygous advantage. Our findings establish a crucial groundwork for future research that utilizes sequencing data from various *P. tricornutum* accessions which we made available via PhaeoEpiView platform (https://PhaeoEpiView.univ-nantes.fr) [10] for easy and comprehensive use. This will enhance our understanding of diatom biology, foster advancements in biotechnology applications, and optimize trait selection.

## Acknowledgements

We thank Carine Pruvost for media preparation. We are grateful to Achal Rastogi for his helpful discussions on the bioinformatics methodology related to phylogeny. LT acknowledges support from the region of Pays de la Loire and Nantes métropole (ConnecTalent EPIALG project), Epicycle ANR project (ANR-19-CE20-0028-02) and Région Pays de la Loire ImpulseAlgae projects. FY was supported by CSC Grant 201906310152. We are grateful to the bioinformatics core facility of Nantes University (BiRD Biogenouest) for its technical support.

## Authors contribution

LT conceived and designed the study. TC performed and coordinated the bioinformatics analysis. FY conducted most of the experiments. AG extracted RNA and performed the PAM study. EM contributed to the bioinformatics analysis. BJ supervised the PAM study and assisted with data analysis. GP and LT supervised the genetic population study. OAM performed the WGCNA analysis. YW and UC assisted with DNA extraction. TC, FY, AG, GP and LT analysed and interpreted the data. LT supervised and coordinated the study. LT wrote the manuscript with input from TC, FY, AG, EM and OAM. All authors read and edited the manuscript.

## Data availability

The data that support the findings of this study are openly available in BioProjects PRJNA430316 and PRJNA971163

## Supplementary figures

**Figure S1**. Light microscopy images of *P. tricornutum* accessions depicting cell morphology and size. The red-framed cells (Pt3 in **a,** and Pt9 in **b**) represent the proportions of oval morphotype. **c** Cell size and morphology of Pt11 (**c**), Pt12 (**d**), Pt13 (**e**), Pt14 (**f**) Pt15 (**g**). **h** Cell size, morphology and proportion of tri-radiate morphotype in Pt16. **i** Cell size and morphology in Pt17. Red lines indicate the 30 cells for which width and length were measured.

**Figure S2**. Gel images of PCR product illustrating the validation of INDELs observed in Pt11 to Pt17. **a** Insertion validation. **b** Deletion validation. The molecular weight marker is 1 kb plus DNA ladder (M), and the negative control is represented by (N).

**Figure S3**. Gel images of PCR product illustrating the validation of gene loss observed in Pt11 to Pt17. The molecular weight marker is 1 kb plus DNA ladder (M), and the negative control is represented by (N).

**Figure S4**. The plot shows the error of ADMIXTURE cross-validation process for K ranging from 1 to 17, from all accessions callable SNPs and INDELs. The lowest value (15) gives an indication of the ancestral populations number.

**Figure S5**. Inter-samples correlation heat map. The correlation coefficient is represented by the square of the Pearson correlation coefficient (R). The greater the value of R, the higher the degree of similarity between samples.

**Figure S6**. Comparative transcriptional analysis of genes involved in non-photochemical quenching (NPQ) capacity across accessions using Pt1 8.6 as reference strain. **a** Light-harvesting complex X1 (Lhcx1). **b** Light-harvesting complex X2 (Lhcx2). **c** Light-harvesting complex X3 (Lhcx3). **d** Light-harvesting complex X3 ((Lhcx4). **e** Zeaxanthin epoxidase 1 (ZEP1). **f** Zeaxanthin epoxidase 2 (ZEP2). **g** Zeaxanthin epoxidase 3 (ZEP3).

**Figure S7.** GO enrichment analysis. **a** GO terms enrichment in each of Pt1, Pt7, Pt9, Pt11, Pt12, Pt16 and Pt17. In other accessions, no significant enrichment of GO terms was found. **b** The enrichment of GO terms was performed on the set of specific downregulated DEGs for each ecotype. Bar charts show which GO terms are represented and in which ecotype (Pt3, Pt8, Pt10, Pt11, Pt14, Pt16, Pt17). In other ecotypes, no significant GO enrichment terms were found.

**Figure S8.** Heatmap visualization of gene expression and corresponding eigengenes across accessions in the 33 modules. The x-axis of both the heatmap and barplot displays the *P. tricornutum* accessions in the following order: Pt1R1, Pt2R1, Pt2R2, Pt3R1, Pt3R2 until Pt17R2. Each row of the heatmap corresponds to genes belonging to the module. Red color denotes overexpression and green underexpression.

**Table S1**. Sampling location, date and morphological features of the 17 accessions of *P. trio-rutum*

**Table S2**. List of primer sequences used in the study

**Table S3**. INDEL loci identified in Pt11 to Pt17 accessions

**Table S4**. Copy number variation (CNV) identified in Pt1 to Pt17 accessions

**Table S5**. Gene loss in Pt1 to Pt17 accessions

**Table S6**. Loss of function (LoF) loci in Pt1 to Pt17 accessions

**Table S7**. Table of the Fst distance between the different populations

**Table S8**. Genes under balancing selection identified in Pt1 to Pt17 accessions

**Table S9**. List of the novel genes identified in Pt1 to Pt17 accessions

**Table S10**. Repeats identified in novel transcripts

**Table S11**. Non coding RNA and other categories of RNA identified in novel transcripts. The table is composed of Sequence Name :the name of the sequence, RNA size: the length of the original transcript, ORF size:the size of the potential ORF within the sequence, Ficket Score:the Fickett score is a linguistic feature that distinguishes protein-coding RNA and ncRNA accord-ing to the combinational effect of nucleotide composition and codon usage bias, Hexamer Score:the hexamer score is calculated using a log-likelihood ratio to measure differential hex-amer usage between coding and non-coding sequences, Coding Probability:the coding proba-bility assigned to each transcript(Human:coding probability(CP)>=0.364 indicates coding se-quence,Mouse:coding probability(CP)>=0.44 indicates coding sequence and Zebrafish:coding probability(CP)>=0.38 indicates coding sequence), Coding Label:marking for each sequence whether it is a coding, non-coding, or unknown coding potential transcript.

**Table S12.** List of accession specific genes up or down regulated and their GO

**Table S13**. List of genes up or down regulated and their GO in all accessions or per clade

**Table S14**. *P. tricornutum* accessions modules after merging and their annotation

**Table S15.** Module composition of identified clusters and their corresponding GO

## References

1. Falkowski PG: Evolution of the nitrogen cycle and its influence on the biological sequestration of CO2 in the ocean. Nature 1997, 387:272–275.

2. Treguer PJ, De La Rocha CL: The world ocean silica cycle. Ann Rev Mar Sci 2013, 5:477–501.

3. Lauritano C AJ, Hansen E, Albrigtsen M, Escalera L, Esposito F, Helland K, Hanssen KØ, Romano G and Ianora A: Bioactivity Screening of Microalgae for Antioxidant, Anti-Inflammatory, Anticancer, Anti-Diabetes, and Antibacterial Activities. Front Mar Sci 2016, 3:68.

4. Rabiee N, Khatami, M., Jamalipour Soufi, G., Fatahi, Y., Iravani, S., & Varma, R. S. : Diatoms with invaluable applications in nanotechnology, biotechnology, and biomedicine: recent advances. ACS Biomaterials Science and Engineering 2021, 7:3053–3068.

5. Falciatore A, Jaubert M, Bouly JP, Bailleul B, Mock T: Diatom Molecular Research Comes of Age: Model Species for Studying Phytoplankton Biology and Diversity. Plant Cell 2020, 32:547–572.

6. Veluchamy A, Rastogi A, Lin X, Lombard B, Murik O, Thomas Y, Dingli F, Rivarola M, Ott S, Liu X, et al: An integrative analysis of post-translational histone modifications in the marine diatom Phaeodactylum tricornutum. Genome Biol 2015, 16:102.

7. Rastogi A, Maheswari U, Dorrell RG, Vieira FRJ, Maumus F, Kustka A, McCarthy J, Allen AE, Kersey P, Bowler C, Tirichine L: Integrative analysis of large scale transcriptome data draws a comprehensive landscape of Phaeodactylum tricornutum genome and evolutionary origin of diatoms. Sci Rep 2018, 8:4834.

8. Rastogi A, Murik O, Bowler C, Tirichine L: PhytoCRISP-Ex: a web-based and stand-alone application to find specific target sequences for CRISPR/CAS editing. BMC Bioinformatics 2016, 17:261.

9. Nymark M, Sharma AK, Sparstad T, Bones AM, Winge P: A CRISPR/Cas9 system adapted for gene editing in marine algae. Sci Rep 2016, 6:24951.

10. Wu Y, Chaumier T, Manirakiza E, Veluchamy A, Tirichine L: PhaeoEpiView: an epigenome browser of the newly assembled genome of the model diatom Phaeodactylum tricornutum. Sci Rep 2023, 13:8320.

11. Rastogi A, Vieira FRJ, Deton-Cabanillas AF, Veluchamy A, Cantrel C, Wang G, Vanormelingen P, Bowler C, Piganeau G, Hu H, Tirichine L: A genomics approach reveals the global genetic polymorphism, structure, and functional diversity of ten accessions of the marine model diatom Phaeodactylum tricornutum. ISME J 2020, 14:347–363.

12. De Martino AM, A. Juan Shi, K.P. Bowler, C.: Genetic and phenotypic characterization of Phaeodactylum tricornutum (Bacillariophyceae) accessions. J Phycol 2007, 43:992–1009.

13. Bailleul B, Rogato A, de Martino A, Coesel S, Cardol P, Bowler C, Falciatore A, Finazzi G: An atypical member of the light-harvesting complex stress-related protein family modulates diatom responses to light. Proc Natl Acad Sci U S A 2010, 107:18214–18219.

14. Abida H, Dolch LJ, Mei C, Villanova V, Conte M, Block MA, Finazzi G, Bastien O, Tirichine L, Bowler C, et al: Membrane glycerolipid remodeling triggered by nitrogen and phosphorus starvation in Phaeodactylum tricornutum. Plant Physiol 2015, 167:118–136.

15. Sprouffske K, Aguilar-Rodriguez J, Wagner A: How Archiving by Freezing Affects the Genome-Scale Diversity of Escherichia coli Populations. Genome Biol Evol 2016, 8:1290–1298.

16. Riesco MF, Robles V: Cryopreservation Causes Genetic and Epigenetic Changes in Zebrafish Genital Ridges. PLoS One 2013, 8:e67614.

17. Wing KM, Phillips MA, Baker AR, Burke MK: Consequences of Cryopreservation in Diverse Natural Isolates of Saccharomyces cerevisiae. Genome Biol Evol 2020, 12:1302–1312.

18. Kram KE, Geiger C, Ismail WM, Lee H, Tang H, Foster PL, Finkel SE: Adaptation of Escherichia coli to Long-Term Serial Passage in Complex Medium: Evidence of Parallel Evolution. mSystems 2017, 2.

19. Russo MT, Aiese Cigliano R, Sanseverino W, Ferrante MI: Assessment of genomic changes in a CRISPR/Cas9 Phaeodactylum tricornutum mutant through whole genome resequencing. PeerJ 2018, 6:e5507.

20. Bulankova P, Sekulic M, Jallet D, Nef C, van Oosterhout C, Delmont TO, Vercauteren I, Osuna-Cruz CM, Vancaester E, Mock T, et al: Mitotic recombination between homologous chromosomes drives genomic diversity in diatoms. Curr Biol 2021, 31:3221–3232 e3229.

21. Hafker NS, Andreatta G, Manzotti A, Falciatore A, Raible F, Tessmar-Raible K: Rhythms and Clocks in Marine Organisms. Ann Rev Mar Sci 2023, 15:509–538.

22. Tirichine L, Bowler C: Decoding algal genomes: tracing back the history of photosynthetic life on Earth. Plant J 2011, 66:45–57.

23. Cruz de Carvalho MH, Sun HX, Bowler C, Chua NH: Noncoding and coding transcriptome responses of a marine diatom to phosphate fluctuations. New Phytol 2016, 210:497–510.

24. Nguyen DT, Wu B, Long H, Zhang N, Patterson C, Simpson S, Morris K, Thomas WK, Lynch M, Hao W: Variable Spontaneous Mutation and Loss of Heterozygosity among Heterozygous Genomes in Yeast. Mol Biol Evol 2020, 37:3118–3130.

25. Vartanian M, Descles J, Quinet M, Douady S, Lopez PJ: Plasticity and robustness of pattern formation in the model diatom Phaeodactylum tricornutum. The New phytologist 2009, 182:429–442.

26. Ralph P.J. GR: Rapid light curves: A powerful tool to assess photosynthetic activity. Aquatic Botany 2005, 82:222–237.

27. Platt T, Gallegos, C.L., Harrison, W.G. : Photoinhibition of photosynthesis in natural assemblages of marine phytoplankton. J Mar Res 1980, 38:687–701.

28. Serodio J, Lavaud J: A model for describing the light response of the nonphotochemical quenching of chlorophyll fluorescence. Photosynth Res 2011, 108:61–76.

29. Richards E RM, Rogers S. : Preparation of genomic DNA from plant tissue. Current protocols in molecular biology 1994, 27:1–23.

30. Nguyen TN, Berzano M, Gualerzi CO, Spurio R: Development of molecular tools for the detection of freshwater diatoms. J Microbiol Methods 2011, 84:33–40.

31. Siaut M, Heijde M, Mangogna M, Montsant A, Coesel S, Allen A, Manfredonia A, Falciatore A, Bowler C: Molecular toolbox for studying diatom biology in Phaeodactylum tricornutum. Gene 2007, 406:23–35.

32. Bolger AM, Lohse M, Usadel B: Trimmomatic: a flexible trimmer for Illumina sequence data. Bioinformatics 2014, 30:2114–2120.

33. Vasimuddin M, Misra, S., Li, H. and Aluru, S.: Efficient Architecture-Aware Acceleration of BWA-MEM for Multicore Systems. In IEEE International Parallel and Distributed Processing Symposium (IPDPS). pp. 314–324. Rio de Janeiro, Brazil; 2019:314–324.

34. van der Auwera G, O’Connor, B.D.: Genomics in the Cloud: Using Docker, GATK, and WDL in Terra. 2020.

35. Cingolani P, Platts A, Wang le L, Coon M, Nguyen T, Wang L, Land SJ, Lu X, Ruden DM: A program for annotating and predicting the effects of single nucleotide polymorphisms, SnpEff: SNPs in the genome of Drosophila melanogaster strain w1118; iso-2; iso-3. Fly (Austin) 2012, 6:80–92.

36. Korneliussen TS, Albrechtsen A, Nielsen R: ANGSD: Analysis of Next Generation Sequencing Data. BMC Bioinformatics 2014, 15:356.

37. R: A language and environment for statistical computing [https://www.R-project.org/.]

38. Alexander DH, Novembre J, Lange K: Fast model-based estimation of ancestry in unrelated individuals. Genome Res 2009, 19:1655–1664.

39. Liao Y, Smyth GK, Shi W: featureCounts: an efficient general purpose program for assigning sequence reads to genomic features. Bioinformatics 2014, 30:923–930.

40. Conway JR, Lex A, Gehlenborg N: UpSetR: an R package for the visualization of intersecting sets and their properties. Bioinformatics 2017, 33:2938–2940.

41. Fijarczyk A, Babik W: Detecting balancing selection in genomes: limits and prospects. Mol Ecol 2015, 24:3529–3545.

42. Galili T: dendextend: an R package for visualizing, adjusting and comparing trees of hierarchical clustering. Bioinformatics 2015, 31:3718–3720.

43. Schliep KP: phangorn: phylogenetic analysis in R. Bioinformatics 2011, 27:592–593.

44. Kim D, Paggi JM, Park C, Bennett C, Salzberg SL: Graph-based genome alignment and genotyping with HISAT2 and HISAT-genotype. Nat Biotechnol 2019, 37:907–915.

45. Ito K, Murphy D: Application of ggplot2 to Pharmacometric Graphics. CPT Pharmacometrics Syst Pharmacol 2013, 2:e79.

46. Pertea M, Pertea GM, Antonescu CM, Chang TC, Mendell JT, Salzberg SL: StringTie enables improved reconstruction of a transcriptome from RNA-seq reads. Nat Biotechnol 2015, 33:290–295.

47. Afgan E, Baker D, Batut B, van den Beek M, Bouvier D, Cech M, Chilton J, Clements D, Coraor N, Gruning BA, et al: The Galaxy platform for accessible, reproducible and collaborative biomedical analyses: 2018 update. Nucleic Acids Res 2018, 46:W537–W544.

48. Langfelder P, Horvath S: WGCNA: an R package for weighted correlation network analysis. BMC Bioinformatics 2008, 9:559.

49. Love MI, Huber W, Anders S: Moderated estimation of fold change and dispersion for RNA-seq data with DESeq2. Genome Biol 2014, 15:550.

50. Ait-Mohamed O, Novak Vanclova AMG, Joli N, Liang Y, Zhao X, Genovesio A, Tirichine L, Bowler C, Dorrell RG: PhaeoNet: A Holistic RNAseq-Based Portrait of Transcriptional Coordination in the Model Diatom Phaeodactylum tricornutum. Front Plant Sci 2020, 11:590949.

51. Zhao X, Rastogi A, Deton Cabanillas AF, Ait Mohamed O, Cantrel C, Lombard B, Murik O, Genovesio A, Bowler C, Bouyer D, et al: Genome wide natural variation of H3K27me3 selectively marks genes predicted to be important for cell differentiation in Phaeodactylum tricornutum. New Phytol 2021, 229:3208–3220.

52. Watterson GA: On the number of segregating sites in genetical models without recombination. Theor Popul Biol 1975, 7:256–276.

53. Siewert KM, Voight BF: Detecting Long-Term Balancing Selection Using Allele Frequency Correlation. Mol Biol Evol 2017, 34:2996–3005.

54. Lepetit B, Sturm S, Rogato A, Gruber A, Sachse M, Falciatore A, Kroth PG, Lavaud J: High light acclimation in the secondary plastids containing diatom Phaeodactylum tricornutum is triggered by the redox state of the plastoquinone pool. Plant Physiol 2013, 161:853–865.

55. Jahns P, Holzwarth AR: The role of the xanthophyll cycle and of lutein in photoprotection of photosystem II. Biochim Biophys Acta 2012, 1817:182–193.

56. Taddei L, Stella GR, Rogato A, Bailleul B, Fortunato AE, Annunziata R, Sanges R, Thaler M, Lepetit B, Lavaud J, et al: Multisignal control of expression of the LHCX protein family in the marine diatom Phaeodactylum tricornutum. J Exp Bot 2016, 67:3939–3951.

57. Malerba ME, Palacios MM, Palacios Delgado YM, Beardall J, Marshall DJ: Cell size, photosynthesis and the package effect: an artificial selection approach. New Phytol 2018, 219:449–461.

58. Spanbauer TL, Fritz, S.C. and Baker, P.A.: Punctuated changes in the morphology of an endemic diatom from Lake Titicaca. Paleobiology 2018, 44:89–100.

59. Pfeifer SP: From next-generation resequencing reads to a high-quality variant data set. Heredity (Edinb) 2017, 118:111–124.

60. Smith NMA, Wade C, Allsopp MH, Harpur BA, Zayed A, Rose SA, Engelstadter J, Chapman NC, Yagound B, Oldroyd BP: Strikingly high levels of heterozygosity despite 20 years of inbreeding in a clonal honey bee. J Evol Biol 2019, 32:144–152.

61. Kardos M, Akesson M, Fountain T, Flagstad O, Liberg O, Olason P, Sand H, Wabakken P, Wikenros C, Ellegren H: Genomic consequences of intensive inbreeding in an isolated wolf population. Nat Ecol Evol 2018, 2:124–131.

62. Guo L, Zhang S, Rubinstein B, Ross E, Alvarado AS: Widespread maintenance of genome heterozygosity in Schmidtea mediterranea. Nat Ecol Evol 2016, 1:19.

63. Behe MJ: Experimental evolution, loss-of-function mutations, and “the first rule of adaptive evolution”. Q Rev Biol 2010, 85:419–445.

64. Caseys C: Loss of Function, a Strategy for Adaptation in Arabidopsis. Plant Cell 2019, 31:935.

65. Hottes AK, Freddolino PL, Khare A, Donnell ZN, Liu JC, Tavazoie S: Bacterial adaptation through loss of function. PLoS Genet 2013, 9:e1003617.

66. Meyer RS, Purugganan MD: Evolution of crop species: genetics of domestication and diversification. Nat Rev Genet 2013, 14:840–852.

67. Murray AW: Can gene-inactivating mutations lead to evolutionary novelty? Curr Biol 2020, 30:R465–R471.

68. Tournamille C, Colin Y, Cartron JP, Le Van Kim C: Disruption of a GATA motif in the Duffy gene promoter abolishes erythroid gene expression in Duffy-negative individuals. Nat Genet 1995, 10:224–228.

69. de Valles-Ibanez G, Hernandez-Rodriguez J, Prado-Martinez J, Luisi P, Marques-Bonet T, Casals F: Genetic Load of Loss-of-Function Polymorphic Variants in Great Apes. Genome Biol Evol 2016, 8:871–877.

70. MacArthur DG, Balasubramanian S, Frankish A, Huang N, Morris J, Walter K, Jostins L, Habegger L, Pickrell JK, Montgomery SB, et al: A systematic survey of loss-of-function variants in human protein-coding genes. Science 2012, 335:823–828.

71. Labboun S, Terce-Laforgue T, Roscher A, Bedu M, Restivo FM, Velanis CN, Skopelitis DS, Moschou PN, Roubelakis-Angelakis KA, Suzuki A, Hirel B: Resolving the role of plant glutamate dehydrogenase. I. In vivo real time nuclear magnetic resonance spectroscopy experiments. Plant Cell Physiol 2009, 50:1761–1773.

72. Grzechowiak M, Sliwiak J, Jaskolski M, Ruszkowski M: Structural Studies of Glutamate Dehydrogenase (Isoform 1) From Arabidopsis thaliana, an Important Enzyme at the Branch-Point Between Carbon and Nitrogen Metabolism. Front Plant Sci 2020, 11:754.

73. Terce-Laforgue T, Clement G, Marchi L, Restivo FM, Lea PJ, Hirel B: Resolving the Role of Plant NAD-Glutamate Dehydrogenase: III. Overexpressing Individually or Simultaneously the Two Enzyme Subunits Under Salt Stress Induces Changes in the Leaf Metabolic Profile and Increases Plant Biomass Production. Plant Cell Physiol 2015, 56:1918–1929.

74. Li S, Shao Z, Lu C, Yao J, Zhou Y, Duan D: Glutamate Dehydrogenase Functions in Glutamic Acid Metabolism and Stress Resistance in Pyropia haitanensis. Molecules 2021, 26.

75. Julenius K, Pedersen AG: Protein evolution is faster outside the cell. Mol Biol Evol 2006, 23:2039–2048.

76. Jancek S, Gourbiere S, Moreau H, Piganeau G: Clues about the genetic basis of adaptation emerge from comparing the proteomes of two Ostreococcus ecotypes (Chlorophyta, Prasinophyceae). Mol Biol Evol 2008, 25:2293–2300.

77. Malaria Genomic Epidemiology N, Band G, Rockett KA, Spencer CC, Kwiatkowski DP: A novel locus of resistance to severe malaria in a region of ancient balancing selection. Nature 2015, 526:253–257.

78. Key FM, Teixeira JC, de Filippo C, Andres AM: Advantageous diversity maintained by balancing selection in humans. Curr Opin Genet Dev 2014, 29:45–51.

79. Segurel L, Thompson EE, Flutre T, Lovstad J, Venkat A, Margulis SW, Moyse J, Ross S, Gamble K, Sella G, et al: The ABO blood group is a trans-species polymorphism in primates. Proc Natl Acad Sci U S A 2012, 109:18493–18498.

80. Junprung W. SP, Tassanakajon A., Van Stappen G., Peter Bossier: Balancing selection at the ATP binding site of heat shock cognate 70 (HSC70) contributes to increased thermotolerance in Artemia franciscana. Acquaculture 2021, 531:735988.

81. Li XH, Babu MM: Human Diseases from Gain-of-Function Mutations in Disordered Protein Regions. Cell 2018, 175:40–42.

82. Meyer Kea: Mutations in disordered regions can cause disease by creating dileucine motifs. Cell 2018, 175:239–253.

83. Jaenisch R, Bird, A.: Epigenetic regulation of gene expression: how the genome integrates intrinsic and environmental signals. Nat Genet 2003, 33:245–254.

84. Veluchamy A, Lin X, Maumus F, Rivarola M, Bhavsar J, Creasy T, O’Brien K, Sengamalay NA, Tallon LJ, Smith AD, et al: Insights into the role of DNA methylation in diatoms by genome-wide profiling in Phaeodactylum tricornutum. Nat Commun 2013, 4.

